# Screening and characterization of aging regulators using synthesized yeast chromosome *XIII*

**DOI:** 10.1101/2023.11.07.566118

**Authors:** Chun Zhou, Yun Wang, Yikun Huang, Yongpan An, Xian Fu, Daqian Yang, Yilin Wang, Jintao Zhang, Leslie A. Mitchell, Joel S. Bader, Yizhi Cai, Junbiao Dai, Jef D. Boeke, Zhiming Cai, Zhengwei Xie, Yue Shen, Weiren Huang

**Affiliations:** Department of Urology, Shenzhen Institute of Translational Medicine, Shenzhen Second People’s Hospital, The First Affiliated Hospital of Shenzhen University, Shenzhen 518035, China; Guangdong Provincial Key Laboratory of Genome Read and Write, BGI Research-Shenzhen, BGI, Shenzhen, 518083, China; Peking University International Cancer Institute, Peking University Health Science Center, Peking University, Beijing, China; Guangdong Provincial Key Laboratory of Synthetic Genomics and Shenzhen Key Laboratory of Synthetic Genomics. Shenzhen Institute of Synthetic Biology, Shenzhen Institute of Advanced Technology, Chinese Academy of Sciences, Shenzhen, China; BGI Research-Changzhou, BGI, Changzhou, 213000, China; International Cancer Center, Shenzhen University School of Medicine, Shenzhen 518052, China; College of Life Sciences, University of Chinese Academy of Sciences, Beijing, 100049, China; Institute for Systems Genetics and Department of Biochemistry and Molecular Pharmacology, NYU Langone Health, New York, NY 10016, USA; Department of Biomedical Engineering, Johns Hopkins University, Baltimore, Maryland, USA; Department of Biomedical Engineering, NYU Tandon School of Engineering, Brooklyn NY 11201; Manchester Institute of Biotechnology, University of Manchester, 131 Princess Street, Manchester, M1 7DN, UK

**Keywords:** *synXIII*, replicative aging, *HSP104*, SCRaMbLE, *RRN9*, aging regulators, *Saccharomyces cerevisiae*

## Abstract

Yeast, a single eukaryotic cell model organism, demonstrates a progressive aging process. In the era of synthetic biology, study of the impact of synthetic chromosomes and aging is urgent and intriguing. Herein, we successfully constructed the 884 Kb *synXIII* of *Saccharomyces cerevisiae* and conducted replicative aging studies using the synthetic strains. We verified that the rRNA-related transcriptional factor *RRN9* is a major positive controller of replicative lifespan. Using SCRaMbLE and an *HSP104* reporter as a biomarker for mutant discovery, we screened 135 SCRaMbLEd synXIII strains with extended lifespan and identified 10 genes on *synXIII* that potentially serve as aging regulators. In addition, the genome-scale regression analysis of long-replicative lifespan SCRaMbLEd strains revealed distinct dysregulation of nucleus, ribosome, and mitochondrion function networks. Our findings suggest that Sc2.0 yeast has potential for unveiling new aging-related genes and gene-gene interactions underlying replicative lifespan.

## Introduction

As a universal trait observed across the evolutionary spectrum, aging is identified as a critical risk factor for many human pathologies, such as Alzheimer’s disease, Parkinson’s disease, tumor progression, and cardiovascular dysfunction.^1^ Aging is also reported in relation to genomic instability, the loss of proteostasis, deregulated nutrient-sensing, epigenetic alterations, telomere attrition, mitochondrial function.^2–5^ Early aging studies focus on identifying aging-related phenotypes.^6^ ^7^ The focus gradually transitioned to investigate genetic networks underlying these phenotypes and revealed a complex network of intracellular signaling pathways and higher-order processes.^8^ However, it is only 40 years since a new era in aging research was inaugurated after the isolation of the first long-lived strains in Caenorhabditis elegans.^9^ In recent years, there has been verified that the rate of aging is at least partly controlled by evolutionarily conserved genetic pathways and biological processes. The complexity of potential interacting networks creates a considerable challenge to aging studies.

The budding yeast, *Saccharomyces cerevisiae*, has emerged as a powerful chemical and genetic screening platform, which has been widely used as a genetic model for exploring mechanism of aging, since a series of conserved genes and pathways between yeast and human. In particular, the replicative aging model, which measures the number of divisions a yeast mother cell can make before entering into a senescent state, is particularly relevant.^10–12^ However, there are still several difficulties in studying aging: 1) The current technique, including knock-out or overexpression screening and *etc.* to identify longevity factors is inefficient and requires massive screening efforts because measuring lifespan is tedious and labor intensive and is typically performed on single clones; 2) There is a lack of molecular biomarkers available to screen potential factors for replicative aging. Thus, careful assessment of the aging process is required to increase the discovery of downstream medical applications.

With the rapid development of DNA synthesis and manipulation techniques, synthetic biology is increasing in scope with respect to the systematic engineering of model organisms and diverse applications.^13–15^

Concurrent developments of scarless nucleic acid assembly,^16^ and the ability to manipulate extensive genes or genome editing and synthesis, have allowed scientists to rapidly updating numerous genetic designs to optimize system function.^17^ As an example, the *Sc2.0* project generated a series of synthetic yeast chromosomes and further applied the synthetic system in obtaining the desired phenotypes by the controlled large-scale chromosomal rearrangements ^18–27^. As a unique feature of Sc2.0, SCRaMbLE (Synthetic Chromosome Rearrangement and Modification by LoxPsym-mediated Evolution) system, which could produce inversions or deletions or duplications until intentional Cre recombinase induction, has been demonstrated as an effective genome-wide rearrangement tool for genetic screening. It could be a promising tool to study biological questions. This reinforced our interest in using synthetic chromosomes to obtain potential aging regulators, biomarkers, even new insights into aging mechanisms.^19,24,28–30^ In this study, we construct Sc2.0 chromosome XIII (*synXIII*) from scratch and used it as a platform for subsequent aging analysis. Our study fortuitously identified a new aging factor gene, *RRN9* during the process of synthetic chromosome assembly. A design feature (synonymous sequence recoding by PCRTag, with recoded sequence unique to either the wild type or synthetic genome) introduced into this gene increased its expression level and consequently induced lifespan extension. Once *synXIII* assembly was completed, we then deployed a *HSP104* based reporter for detecting strains with extended lifespan. We identified 135 SCRaMbLEd yeast candidates with various chromosomal rearrangements limited to *synXIII*. Then, whole-genome sequencing, combined with deployment of a microfluidic device and time-lapse microscopy-based assay for replicative aging, revealed 10 genes that might play important roles in lifespan changes. Global transcriptome analysis of 20 long-lived strains out of the 135 uncovered 5 longevity genes involved in cell cycle, cytoplasmic translation, mitochondrion organization at an early replicative age are well correlated with lifespan. Our study demonstrates that the Sc2.0 yeast and its built-in SCRaMbLE system could serve as a unique model system for studying aging.

## Results

### Construction and characterization of synXIII

Based on the Sc2.0 project principles,^14^ we designed and constructed the 883,749 bp synthetic chromosome XIII (*synXIII*) (**Figure 1A**, **Table S1**), containing 333 loxPsym sites insertion (loxPsym sequences are 34-bp nondirectional loxP sites capable of recombining in either orientation) and 68 deletions, a total of 100 TAG to TAA conversions, and 629 synthetic PCRtag pairs. Overall, based on the design, *synXIII* is condensed by 4.4% in comparison with its native sequence.

**Figure 1.**
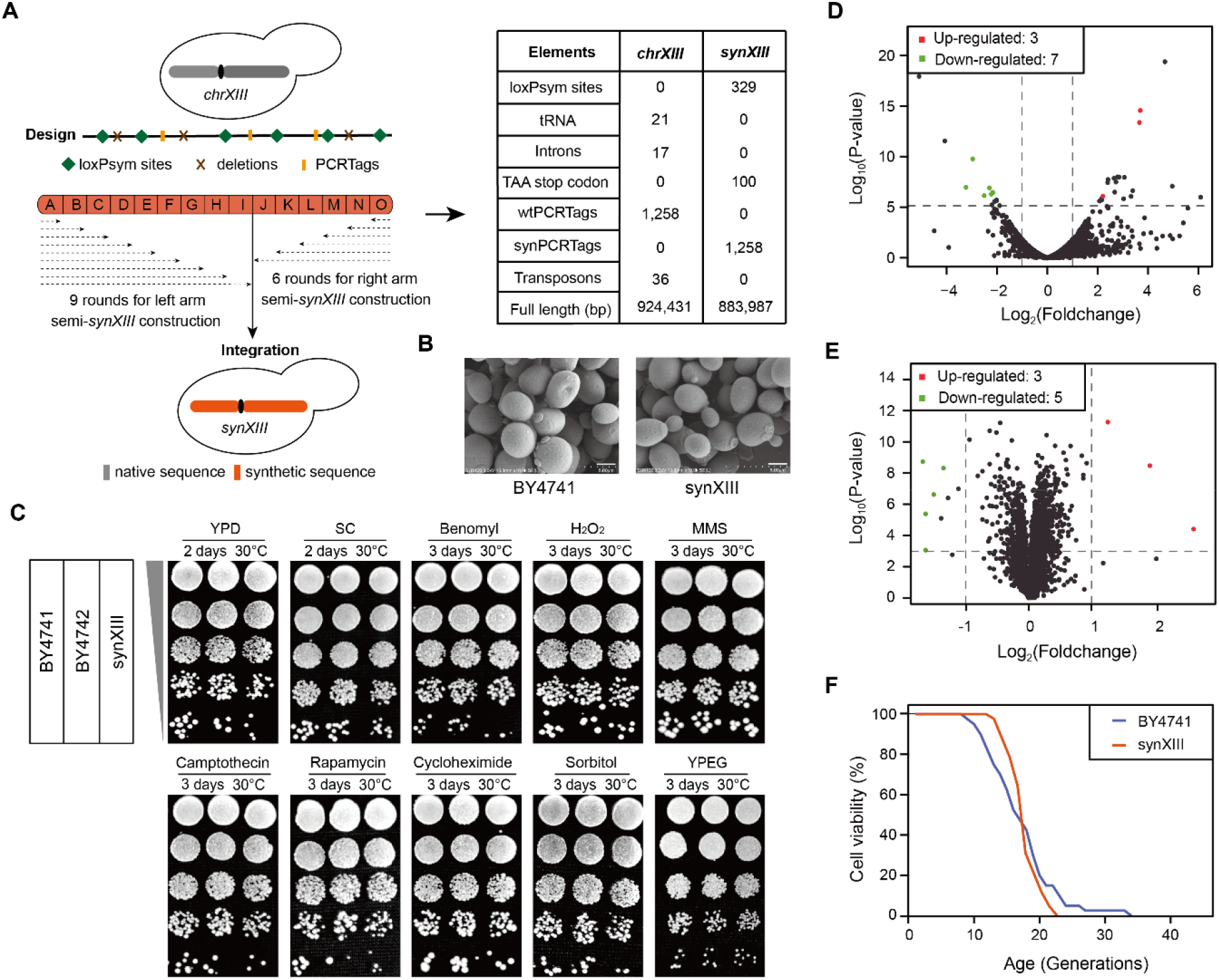
Design, construction, and characterization of synXIII. **(A)** Overview of *synXIII* construction with the indicated designer features. The dotted line represents the range of the synthesized area after each round of replacement, and the arrow represents the direction of integration; The right table listed the difference between the *synXIII* and *chrXIII*. **(B)** Scanning Electron Microscopy (SEM) images of BY4741 and synXIII strains. Scale bars, 5.00 um. **(C)** Cellular viability responses of synXIII upon exposure to normal and stress conditions. 10-fold serial dilutions of overnight cultures of synXIII and wild-type (BY4741 and BY4742) strains were used for plating. Conditions include: YPD at 30°C; SC at 30°C; YPD + Benomyl; YPD + H_2_O_2_ (1 mM, 2 h pretreatment); YPD + methyl methane sulfone (MMS, 0.01% v/v); YPD + Camptothecin (0.1 μg/mL); YPD + Rapamycin (0.2 μg/mL); YPD + Cycloheximide (10 μg/ml, 2 h pretreatment); YPD + Sorbitol (1 M); YPD, yeast extract peptone dextrose; YPEG, yeast extract peptone glycerol ethanol; SC, synthetic complete. **(D)** Identified dysregulated genes of synXIII strain at the transcriptome level compared with BY4741. The differentially expressed genes were assessed at genome-wide significance P < 7.56 × 10^−6^ for 5% family-wise error rate based on 6607 genes with at least one mapped read and also corresponding to log_2_ (foldchange) >= 1 or <= −1. **(E)** Identified dysregulated genetic features of synXIII strain at the proteome level compared with BY474. In D and E, the up and down-regulated genes are labeled in red and green respectively. **(F)** Survive curves of synXIII and BY4741. X-axis presents the number of daughter cells generated from the mother cell. Each strain was measured with three replicates and 40 cells were monitored for each replicate.

According to design principle, all tRNA genes on chromosome *XIII* would relocate into the tRNA neochromosome.^17^ However, one tRNA gene, *tQ(CUG)M*, is essential during the construction (**Figure S1A**). Thus, we relocated it to *chrVI* at this stage (**Figure S1B**) and it could be eliminated once the synXIII strain is consolidated into the final synthetic yeast strain carrying tRNA neochromosome.

To construct the *synXIII* rapidly, we adopt hierarchical assembly strategy.^22^ The whole synthetic chromosome XIII was divided into 15 megachunks and each megachunk (up to 70 kb) further splited into chunks (~15kb) and minichunks (~3kb) (**Figure S2A**). The minichunks were assembled to chunks in *vitro* using Gibson assembly strategy.^16^ The replacements were performed from both ends *in vivo*—along with two wild-type strains (BY4741 and BY4742) in parallel. 5’6 chunks as a megachunk replaced the native one using the standard SwAP-In method. ^31^ 9 / 6 rounds (A-I / J-O) transformation were carried out in BY4741/BY4742 to generate left / right semi-synthetic strains of synXIII-I / J, and subsequently integrated into a whole synthetic chromosome XIII (strain ID: Ycz019) based on a *I*-*Sce*I mediated chromosome integration strategy (**Figure 1A**, **Figure S2B**). The *synXIII* was finally confirmed by whole genome sequencing (WGS) and revealed only “patchwork” variations produced by homologous recombination (**Figure S2C and Table S2**), as reported in other finished synthetic chromosomes. ^22,32^

After the synthesis and correction of *synXIII*, we obtained an almost “Perfect” synXIII (strain ID: Ycz020) (**Figure S2C**). We then carried out a series of phenotypic analyses to validate its fitness. The synXIII and native strain are visually indistinguishable under a scanning electron microscope (**Figure 1B**). Growth assays in diverse conditions showed that the synXIII strain exhibited the same phenotype under various conditions compared with the native BY4741 and BY4742 strains (**Figure 1C**). To analyze the cellular and molecular physiology of the synXIII strain, we explored the potential effect from the synthetic chromosome on the transcriptome and proteome. Transcriptome profiling revealed only 10 differentially expressed genes (DEGs) out of 6,606, comprising three genes up-regulated and seven genes down-regulated (**Figure 1D**). Proteomics analysis identified only eight differentially expressed proteins in synXIII compared to the native strain (**Figure 1E**). By employing a U-shape, high-pressure microfluidic chip previously developed in our lab (**Figure S4A**), we measured the replicative lifespan of synXIII in comparison with BY4741 and revealed a similar survival curve between these two strains (**Figure 1F**). Our results demonstrate that *synXIII* has minimal effect on phenotypic fitness and cell physiology, it displays a replicative lifespan that is comparable to that of the wild-type strain.

### Elevated *RRN9* extended yeast lifespan

The ability of genetic manipulation is very important since the understanding the function or structure of nucleic acid and protein. The designers of synthetic chromosome provide a potential avenue to understand the gene function.^27^ During the progressive construction of synXIII, we observed an interesting finding that the intermediate strain synXIII-N shows an extended lifespan than its parental intermediate stain synXIII-O (**Figure 2A**). By individually integrating each chunk of megachunk N into synXIII-O for lifespan measurement, we found that the synthetic N2 chunk integration strain showed increased lifespan relative to synXIII-O. Then we further identified the modifications introduced in the *RRN9* gene responsible for the increased lifespan (**Figure 2B**).

**Figure 2.**
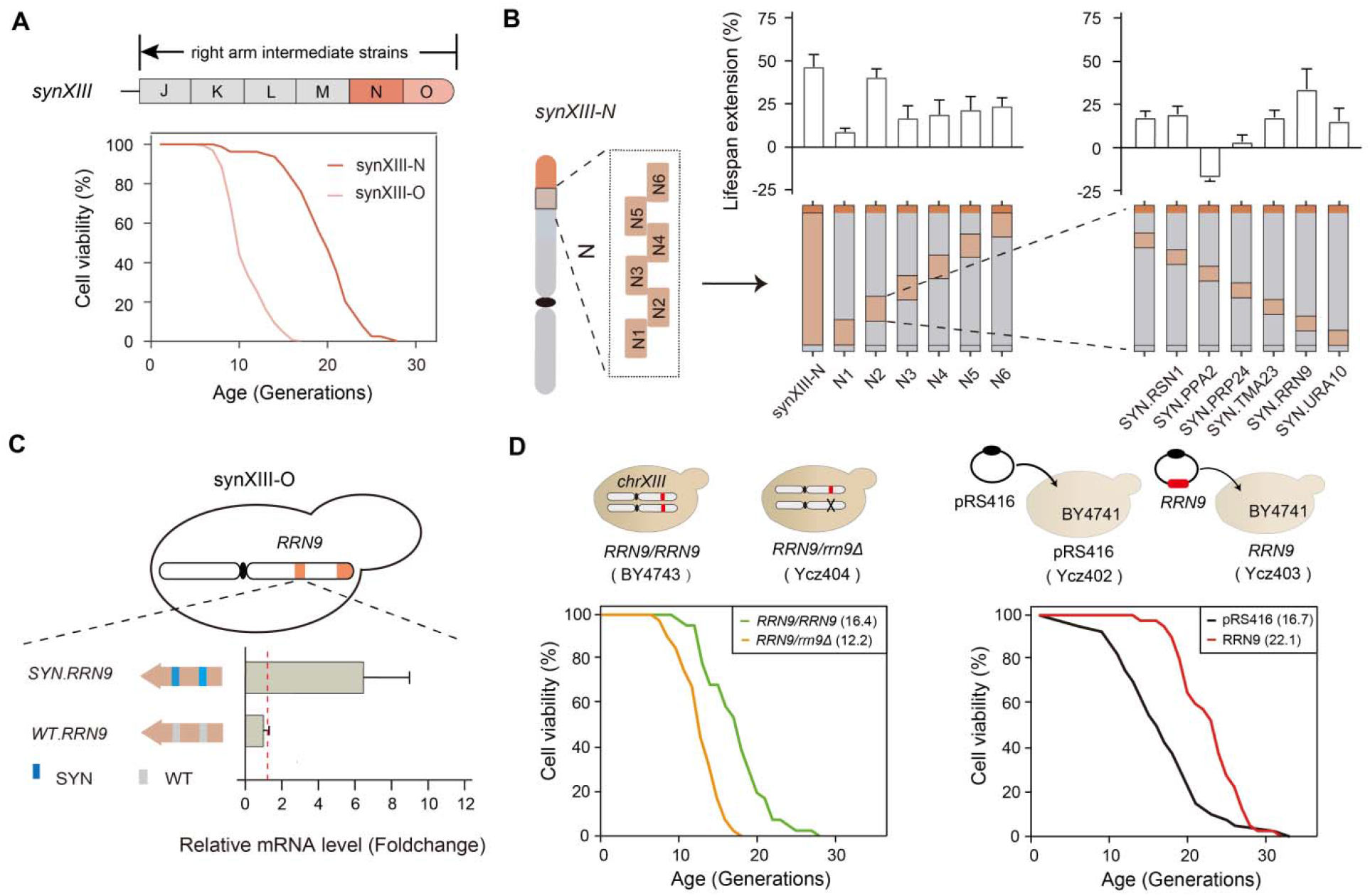
Synthetic *RRN9* exhibits higher expression and promotes lifespan extension. **(A)** Lifespan measurement of intermediate strains, synXIII-O and synXIII-N. The arrow presents the direction of construction. **(B)** The lifespan measurement of all intermediate strains of synthetic megachunk N (highlight in orange) compared to synXIII-O. Synthetic megachunk N was segmented into N1~N6 chunks, which was integrated respectively using synXIIIO as the host strain. N2 chunk carried six synthetic genes (*RSN2, PPA2, PRP24, TMA23, RRN9, URA10*) and using as the host strains, six genes was integrated respectively in synXIII-O strain. The error bar represents SD from three independent replicates. **(C)** The expression level of synthetic and native *RRN9* gene measured by RT-PCR. The red dashed line represents the expression level of native *RRN9* gene in BY4741. The error bars indicated SD from three independent experiments. The blue and gray bars present the synthetic and native PCRTags respectively. **(D)** Survival curves of the strains with diverse doses of *RRN9*. The Ycz402 and Ycz403 strains express pRS416 (centromere-based empty vector) and pRS416-*RRN9* in BY4741 respectively. Ycz404 is a diploid strain derived from BY4743 with one copy of *RRN9*. The number in parentheses indicates the median generation of the corresponding survival curve. For panel A, B and D, each lifespan measurement was performed with three replicates and 40 cells were monitored for each replicate. The statistical confidence is calculated by one-sided T-test, and error bars represent standard division.

In synXIII, *RRN9* is synonymously recoded with two synthetic PCRtag 1 & 2 (**Figure 2C**). The synthetic PCRTag has been reported to occasionally cause variations in expression levels and growth defects.^33^ Thus, we introduced both synthetic and native version of *RRN9* gene into the synXIII-O strain for RT-PCR analysis. We found that *synRRN9* was up-regulated compared to native *RRN9* in the resulting strains (**Figure 2C**). To our knowledge, whether *RRN9* regulates aging has not been reported. So, we examined the effect of different doses of *RRN9* upon lifespan. First, we made the diploid *RRN9/rrn9*Δ strain to achieve 50% reduction in *RRN9* copy number and measured its lifespan. We found that a *RRN9*/*rrn9*Δ strain exhibited a shorter lifespan compared to the control *RRN9*/*RRN9* strain (**Figure 2D**). Then, we increased the copy number of *RRN9* in the haploid BY4741 strain by introducing extra copy of *RRN9* gene on a centromeric (medium copy number) plasmid and found that the replicative lifespan was significantly increased by 37.5% (**Figure 2D**, P-value <0. 0001).

To determine which PCRTag was responsible for the increased mRNA abundance of *RRN9*, we individually introduced the two PCRtags to synXIII-O strain. We found that the insertion of SYNtag2 was sufficient to cause up-regulation of *RRN9* in synXIII-O (**Figure 3A**), a result that was further confirmed by immunoblotting. The Rrn9 protein abundance of synXIII-O strain with SYNtag2 is increased by ~20% in comparison with that of synXIII-O strain with WTtag2 (**Figure 3B**).

**Figure 3.**
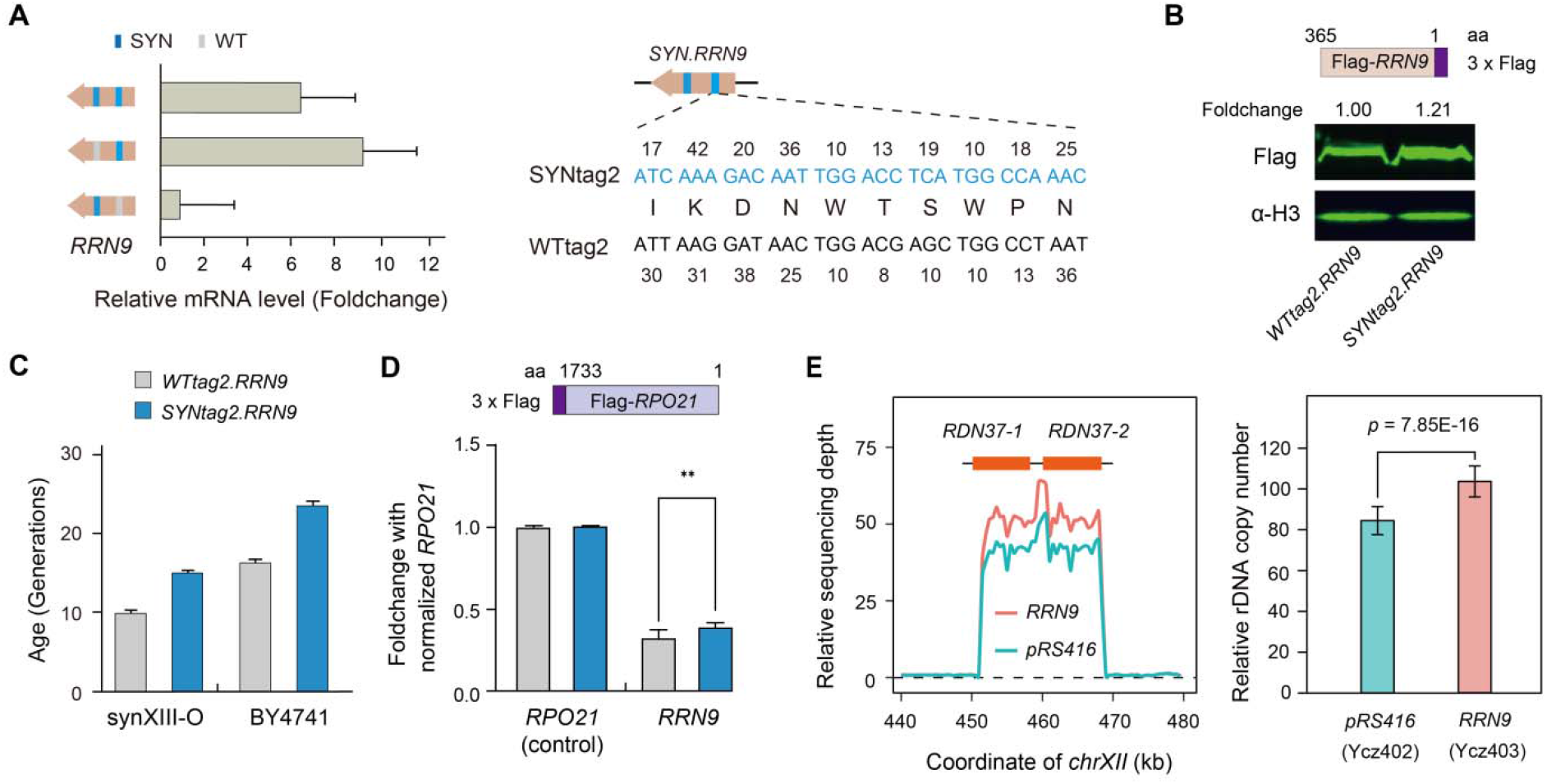
The overexpression of *RRN9* triggers the increase of rDNA copy number and potentially results in an extended lifespan. **(A)** Dissection of the effect of synonymous recoding by PCRTags in *RRN9* gene to its gene expression measured by RT-PCR. The error bars indicate SD from three independent replicates. The synthetic and wildtype PCRTags are presented in blue and gray respectively. The right panel presents the corresponding sequence of synthetic and wildtype PCRtags. **(B)** The corresponding expression level of Rrn9 protein with wildtype and synthetic PCRTag2 by immunoblot. WTtag2 RRN9, expressed from the locus RRN9 in BY4741; SYNTag2 RRN9, expressed from the locus RRN9 with the synthetic PCRtag2 replacing the native PCRtag in BY4741; Protein level was quantified as a fold change. (**C**) The replicative aging measurement of synXIII-O and BY4741 strains with wildtype and synthetic PCRTag2. Error bars indicate SD from three independent replicates. **(D)** RNAPII-ChIP analyses at *RRN9* by RT-PCR. The fold change of RNAPII ratios was normalized to levels of *RPO21* with 3 biological replicates and repeated 3 times. The statistical confidence is calculated by one-sided T-test, and error bars represent SD. **(E)** Estimation of the corresponding rDNA copy number of Ycz403 and Ycz402 by relative sequencing depth. The relative sequencing depth indicated an average of sequencing depth in each 500-bp window normalized to the sequencing depth of whole genome. The statistical confidence is calculated by one-sided T-test, and error bars represent standard division.

In order to confirm the solo effect of SYNtag2 on replicative lifespan, we measured the synXIII-O and BY4741 strains with SYNtag2. Both strains had significantly increased lifespan as compared to the corresponding parental strains (**Figure 3C**), suggesting that the SYNtag2 is responsible for the observed lifespan extension. Next, we probed the potential mechanism for the observed replicative lifespan extension. We examined the RNAPII occupancy by ChIP-qPCR ^27,34^ in the strain carrying SYNtag2 in comparison with the strain carrying WTtag2. Our result showed an increased RNAPII occupancy in *RRN9* locus (**Figure 3D**). *RRN9* encodes a component of the UAF (upstream activating factor) complex, an essential transcription factor for RNA polymerase I,^35,36^ UAF is a major determinant of rDNA copy number in yeast.^37,38^ Then we quantified the rDNA copy number of Ycz403 and Ycz402 by whole genome sequencing and observed that the Ycz403 strain indeed increased rDNA copy number by 22.7% compared to Ycz402 (**Figure 3E**), also discovered the increase by 65% in synXIII-N to synXIII-O (**Figure S4**). We also observed up-regulated expression of *SIR2* in Ycz403 on both the transcriptome and proteome level (**Figure S3A and S3B**). Taken together, we identified the rRNA-related transcriptional factor *RRN9* as a major new positive regulator of replicative lifespan.

**Figure 4.**
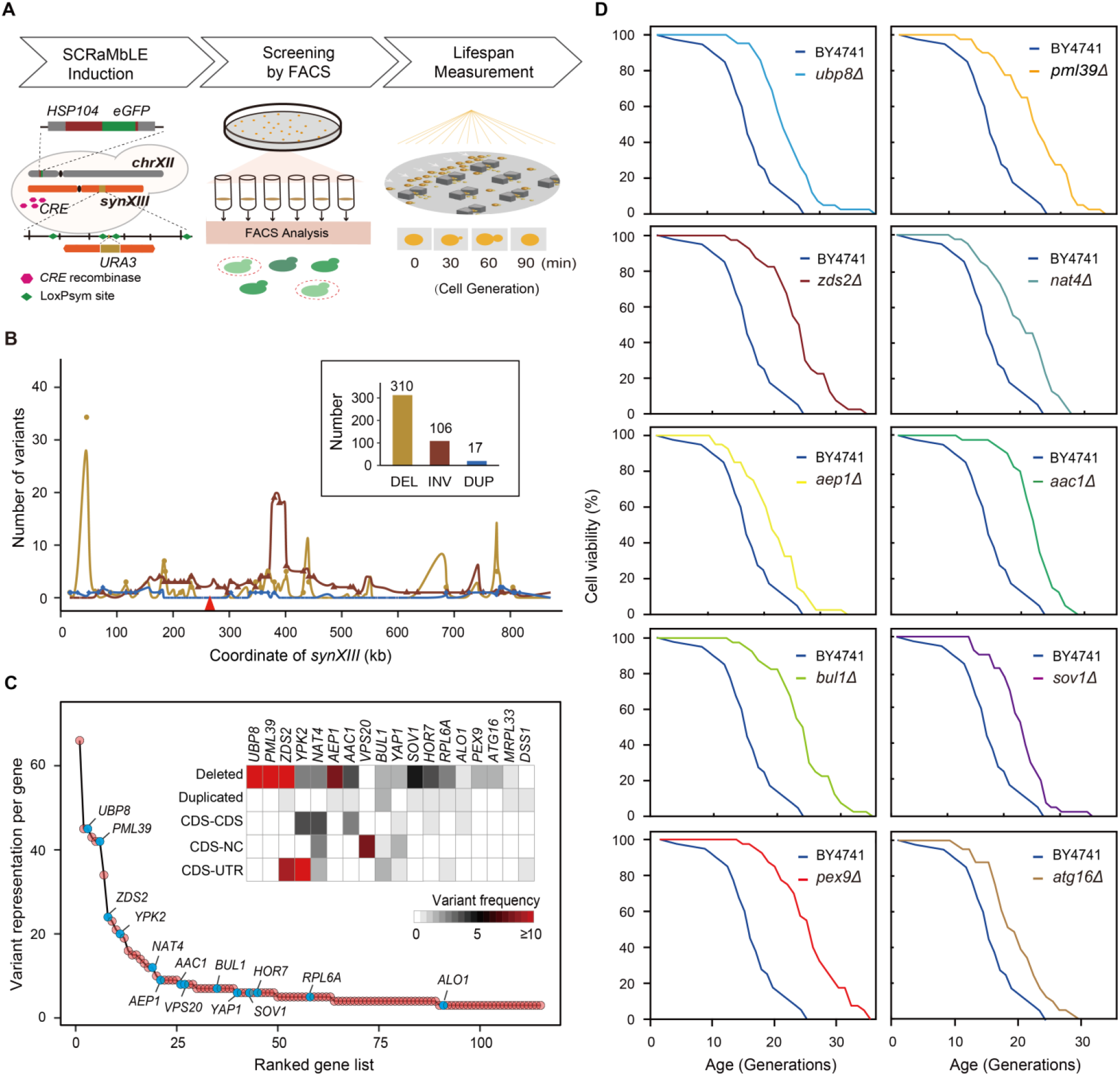
Screening and identification of potential aging regulators from the long-lived SCRaMbLEd synXIII strains. **(A)** Schematic illustration of SCRaMbLE and screening process for long replicative aging strains. The *HSP104* fused to e*GFP* (*HSP104-*e*GFP*) acts as a reporter for replicative lifespan in yeast. The *URA3* reporter was randomly integrated into regions positioned between two loxPsym sites allowing for SCRaMbLEant selection by 5-FOA. SCRaMbLEants with lower fluorescence intensity were selected by FACS, followed by lifespan measurement using a microfluidic device. **(B)** The landscape of SCRaMbLE events including deletion (DEL), inversion (INV), and duplication (DUP) on *synXIII* chromosome of in total 135 screened strains. The red triangle indicated the locus of URA3 insertion on *synXIII*, and the variation representation number of *URA3* deletion was not included in the figure to avoid biased analysis. **(C)** The variant representation per gene (the number of times a variant is represented) in the collection of 135 long-lived SCRaMbLEants. The blue dots represent the selected genes for lifespan measurement in panel D (frequency ≥ 3). The heat map shows the distribution of 5 different variant types for each selected gene. The color indicates the frequency of different rearrangement event types for target genes in these SCRaMbLEd strains. CDS-CDS, CDS-NC, CDS-UTR represent the gene of 3’-UTR with a replacement by a CDS, non-coding (NC) sequence or non-native UTR respectively. **(D)** Survival curves of BY4741 and each single null mutant strains with three replicates (n = 40 for each replicate).

### Generation of a SCRaMbLEant library

Based on a well-constructed *synXIII* with integrated SCRaMbLE system, we sought to screen lifespan-extended-strains from SCRaMbLEd derivatives (SCRaMbLEants) of synXIII first and subsequently decipher their aging regulators, even mechanistic basis. Previously, *HSP104-eGFP* was used as a lifespan reporter within single cells with the same genetic background.^39^ To verify whether the *HSP104-eGFP* reporter is applicable across various yeast mutants, we measured both lifespan and *HSP104-eGFP* level in a series of well-studied mutants, including the well-known short-lived yeast (*ted1*Δ)^40^ and long-lived yeast (*fob1*Δ)^41^ as controls (**Figure S5B**). We observed a strong negative correlation between lifespan and the *HSP104-eGFP* level (Pearson correlation R = −0.916, *p* = 3.986E-06, **Figure S5C**), suggesting that the *HSP104-eGFP* reporter system could be used to screen for long-lived strains. In addition, an auxotrophic marker, *URA3* was integrated into the middle of two loxPsym sites between *YMR011W* and *YMR012W* on *synXIII* (270,117 ~ 270,525 bp) to ensure a powerful positive selection for SCRaMbLEants in the library.^28^ Based on the above-validated lifespan reporter system, we developed a new approach, *HSP104* was fused to an enhanced Green Fluorescent Protein gene (*HSP104-eGFP*) in synXIII strain, generating a reporter for replicative lifespan to achieve high-throughput and effective screening, coupling SCRaMbLE and reduced *HSP104-eGFP* activity to screen for long-lived strains (**Figure 4A**). A pool of about 6000 SCRaMbLEants was obtained and we identified 135 long-lived SCRaMbLEants showing *HSP104-eGFP* levels decreased by at least 10% in comparison with the parental synXIII strain (**Figure S5D**).

### Screen of long-lived SCRaMbLEants yields 10 aging regulator genes

To validate the effectiveness of the above system for aging regulators screening, whole genome sequencing (WGS) of the 135 selected strains was performed to identify potentially causative SCRaMbLEants. A total of 433 recombination events including deletion, inversion and duplication were observed at frequencies of 71.6% (310), 24.5% (106) and 3.9% (17) respectively among these strains (**Figure 4B**). All SCRaMbLE events encompassed 321 genes at varying frequencies, representing ~65% of all genes on *synXIII* (**Table S3**). Five types of gene variations were generated including deleted, duplicated, or 3’-UTR changed (including CDS-CDS, CDS-NC and CDS-Non, Native_UTR). In a 3’-UTR changed gene, the coding domain sequence was joined to a convergent CDS, noncognate UTR (Non_Native_UTR), or other noncoding sequence (NC). We observed 162, 209 and 149 variations with gene deleted, duplicated, or 3’-UTR changed, respectively, that might potentially alter gene expression level and are potentially associated with long-lived phenotypes.

Among the mutated genes, we examined 18 potential lifespan modulatory genes with high variation frequencies (**Figure 4C**), and concluded that deletion of *PML39, ZDS2, UBP8, BUL1, AAC1, ATG16, AEP1, SOV1, NAT4*, and *PEX9* significantly extended lifespan ranging from 20% to 50% (**Figure 4D**), eight out of the ten genes are consistent with previous reports. The genes, *ZDS2,UBP8, BUL1, AEP1, SOV1, NAT4* and *PEX9* were reported as lifespan-extension genes involving functions of chromatin remodeling and activity of mitochondrion and proteasome.^42–47^ *PML39* was reportedly involved in the retention of improper mRNPs in the nucleus and the mutant partially rescued thermosensitive phenotypes of messenger ribonucleoparticle (mRNP) assembly mutants and the mutant show lifespan-extension. ^48,49^ In contrast, *AAC1* and *ATG16* are involved in the mitochondrial inner membrane ADP/ATP translocator and a component of autophagy, respectively.^50,51^ These two genes show potential roles in lifespan extension not reported previously.

### Transcriptome analysis of 20 long-lived strains reveals distinct dysregulation of aging gene networks

To further deepen the application of synthetic chromosome for aging study, we performed transcriptome analyses for 20 long-lived strains with replicative lifespan extensions of 20-50% from the 135 strains collection (**Figure 5A**, **S6A and Table S4**). By applying hierarchical clustering based on the significance of functional enrichment, we found that the 20 strains fell into two groups, which correlate with varying degrees of lifespan extension (Wilcoxon rank sum exact test, *P*-value = 0.03256, **Figure 5B**). The results suggested that replicative lifespan was highly associated with transcriptional level physiological changes. The profiling showed significant up-regulation in the related functions for ribosomes (GO:0005840) and the nucleolus (GO:0002181), and the biological processes of cytoplasmic translation (GO:0002181), and rRNA processing (GO:0006364) (**Figure S6B and S6C**). These up-regulation of above function also observed in the single knock-out mutants in BY4741 (**Figure S6D**). These results suggested that the long-lived strains displayed enhanced capacity for global protein synthesis. ^52^

**Figure 5.**
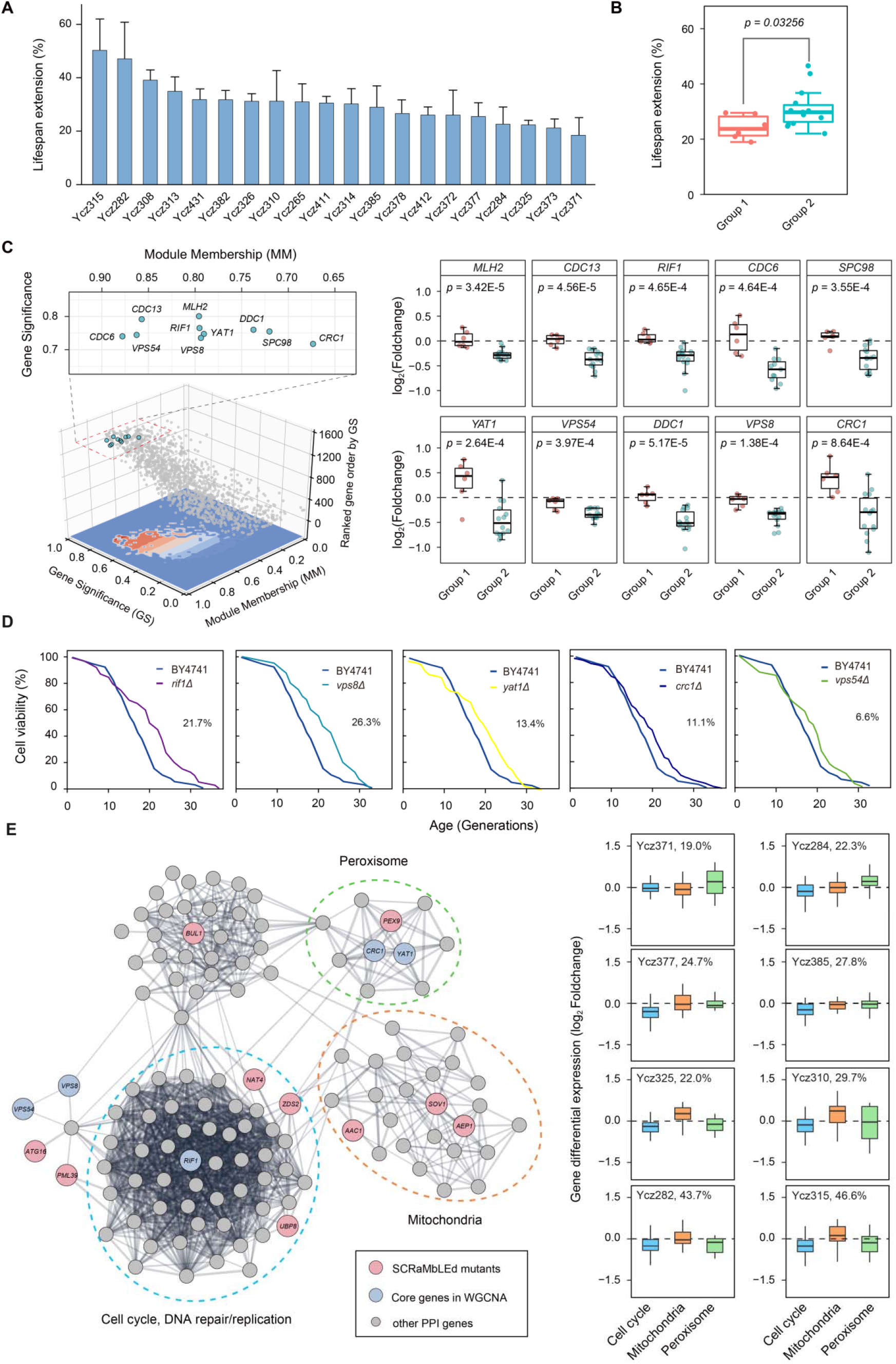
Genome-scale analysis of potential regulatory networks related to aging. **(A)** Lifespan extension of 20 long-lived SCRaMbLEd strains compared to parental strain synXIII Ycz063. Data are presented as the average of value (n = 3). The error bar indicates the SD. **(B)** The cluster analysis of the selected 20 strains in consideration of the replicative lifespan and transcriptome profiling (shown in Figure S5C). The statistical significance was calculated by Wilcoxon rank sum test. **(C)** Screening aging regulators by gene significance (GS, correlation of its gene expression profile with replicative aging) and module membership (MM, correlation of its gene expression profile with the module eigengene). In modules related to a trait of interest genes with high module membership often also have high gene significance. The blue dots indicated the selected aging regulators with GS > 0.7, MM > 0.65 and log_10_MCC > 8. The Maximal Clique Centrality (MCC) was calculated by CytoHubba APP in Cytoscape. Right side: the comparison of differential expression of each gene between the two groups. The p-value was calculated by single-sided T_test. (**D**) Survival curves of BY4741 and each single null mutant strains with three replicates (n = 40 for each replicate). The percentage presents the increase of lifespan for each strain. **(E)** Analysis of protein-protein interaction (PPI) networks that potentially related to aging. The node indicates genes and the width of edge presents the confidence of protein-protein interactions. The network layout was calculated by Edge-weighted Spring Embedded Layout Algorithm in Cytoscape. The dot line indicates each sub-network. The biological function of sub-network was identified by STRING enrichment. Differential gene expression of each sub-network in the lifespan-extended strains. The percentage on the right side of strain id presents the increase of lifespan for each strain.

To gain more insights into the synergistic effects of long-lived mutants, we adopt weighted gene co-expressing network analysis (WGCNA) to identify gene modules. The WGCNA has been proven to be an effective tool to dissect gene modules related to complex phenotypes.^53–56^ By exploiting the normalized transcripts per million (TPM) for each gene in groups 1 and 2, we further identified ten core genes in this module defined by gene significance (GS) and module membership (MM) scores (**Figure 5C left panel and Table S5**), among which, *CDC13*, *RIF1*, *DDC1*, *YAT1*, *VPS54*, *VPS8*, and *CRC1* were consistent with previous reports associated with replicative lifespan.^57–65^ The expression pattern of the genes in group 2 showed significant down-regulated compared to group 1 (**Figure 5C right panel**), which suggests that the decreased expression of these genes might have an important effect on lifespan extension. Thus, we conducted aging measurements on individual null mutants of genes *RIF1*, *VPS8*, *YAT1*, *CRC1*, *VPS54*, *MLH2*, and *DDC1* in BY4741, as well as essential genes *CDC6*, *CDC13*, and *SPC98* in BY4743. Our results showed that deletion of *RIF1*, *VPS8*, *YAT1*, *CRC1*, and *VPS54* genes led to a significant extension of lifespan, ranging from 6.6% to 26.3% (**Figure 5D**).

In order to further comprehend the mechanisms of lifespan, a network of protein-protein interactions (PPI) was established to summarize the functional connection among those aging regulators. The results indicated that these aging regulators were involved in cell cycle, the functions of autophagy, peroxisome, and mitochondria (**Figure 5E left panel**). It has been reported that cell division, reactive oxygen species (ROS) removal, and low energy consumption are associated with replicative lifespan.^66–68^ Our result suggests that in addition to genes on *synXIII* that can directly play a role in the lifespan extension, SCRaMbLE can also introduce chromosomal variations that could lead to lifespan extension by causing synergistic changes in the physiology of cells.

In order to comprehend the interaction of aging gene network effect on lifespan extension, we further dissected these functional changes in the 8 strains out of the above 20 longevity strains. The results exhibited dysregulation patterns of the involved aging gene networks did vary substantially from strain to strain. The gene differential expression of decreased nucleus activity related to cell cycle, DNA repair and replication and increased peroxisome activity show the prominent effect on these strains., whereas increased activity of mitochondria potentially effect inversely(**Figure 5E right panel**). Taken together, our results showed that the aging gene networks might play a combinatorial role in replicative lifespan extension. It will be of great interest to further dissect the interaction of those aging network involved in these aging gene-dependent replicative lifespan.

## Discussion

We developed a new approach based on SCRaMbLE and reduced *HSP104-eGFP* activity for screening the long-lived strains. With the approach, we identified 135 SCRaMbLEant candidates for a study of replicative aging, which is not an intuitively visible phenotype, distinct from color or colony size etc. We produced a library of SCRaMbLEants, deploying an *HSP104-eGFP* aging-associated reporter to enable high-throughput screening for long-lived SCRaMbLEants. As *HSP104-eGFP* level is negatively correlation with lifespan (**Fig. S4C**), and its abundance shows a dynamic increase in the cell during cell divisions.^39^ The long-lived SCRaMbLEd cells with low fluorescence of HSP104-eGFP would underlies in nascent progeny cells in the population, resulting in false positives. Therefore, the screening system requires an accurate measurement fluorescence of populations derived from independent colonies by FACS, rather than by simple single cell sorting.

Our efforts successfully identified multiple targets that can contribute to the yeast lifespan extension. We demonstrated the feasibility and efficiency of this approach for large-scale lifespan factor screening. Specifically, we identified *RRN9* as a new lifespan regulator. We showed that recoding a single PCRTag sequence of 10 codons from *synXIII* led to substantial increases in expression level of *RRN9* and consequent increase in replicative lifespan by 30-45%. *RRN9* is necessary for upstream activation factor (UAF) complex integrity, which is a multifunctional transcription factor in *Saccharomyces cerevisiae* that plays dual roles in activating RNA polymerase I (Pol I) transcription and repression of RNA polymerase II (Pol II).^69^ Decreased expression of *RRN9* activates Pol II. while overexpression of *RRN9* represses it^70^. Therefore, we suggest that *RRN9* positively contributes to lifespan extension by increasing rRNA/ribosome biosynthesis. Meanwhile, we observed a significant increase of rDNA copy number activated by the overexpression of *RRN9*, suggesting that the regulation of ribosomal DNA copy number positively correlated with replicative lifespan as previous study^71^.

Additionally, we also found that reduction of *PML39, ZDS2, UBP8, BUL1, AAC1, ATG16, AEP1, SOV1, NAT4,* and *PEX9* expression also contributed to yeast lifespan extension. Among the 18 genes with high variant representation that we selected, 10 out of 18 null mutants showed a positive correlation with lifespan extension. The results revealed that the method adopted here effectively identified aging genes. Furthermore, we investigated two of the SCRaMbLEants in detail, strain Ycz315, which showed the longest lifespan extension of 46.6% contained two deletions, three inversions and one duplication. On the other hand, strain Ycz411 with a lifespan extension of 29.2% lifespan extension, had two deletions and four inversions. Both strains contain the *PML39* deletion and *AAC1* inversion, but they exhibited lifespan extension values. This result suggests that further gene-gene interactions may contribute to lifespan extension, and that SCRaMbLE may aid in identifying such interaction. By leveraging the 20 long-lived strains, we analyzed the transcriptional perturbation of each long-lived strain combined with the WGCNA method to uncover and verify aging-related genes. This provides an additional avenue for studying aging by using synthetic chromosomes.

In brief, our study revealed that intermediate synthetic chromosome XIII strain, synXIII-N, was useful in its surprising longer than expected lifespan, which was mapped to recoding of just 10 codons in the *RRN9* gene. Previous investigation towards the identification of new aging genes was mainly based on gene knockout or individual overexpression. However, this approach is inefficient considering the potential involvement of multiple genes in the regulatory network. The SCRaMbLE system of Sc2.0 offered us the opportunity to shuffle multiple genes on chromosomal level and reveal potential long and short-lived mutants, as well as genetic interactions involving multiple target genes. Therefore, this approach could serve as an effective way to identify aging-related factors, which hold great potential value for the development of anti-aging drugs and aging-related diseases, as well as aging biomarkers.

### Limitations of the study

There are still some present limitations and potential challenges of this study to be addressed in the future. Firstly, *HSP104* is directly associated with protein aggregation in the cell. By use of *HSP104-eGFP* as a reporter may result in the loss of regulators of aging that are caused by other factors. A more universal and efficient reporting system would be highly desirable to identify long-lived strains with more diverse genotypes relative to replicative aging. Secondly, for 315 SCRaMbLEd synXIII long-lived strains with different genotypes, the underlying mechanisms of lifespan extension needs further in-depth and intensive investigation. Thirdly, as the completion of synthetic Sc 2.0 chromosomes, the involvement of all genes in whole genome would bring challenges in the dissection of the potential correlation between genotypes and aging-related phenotypes.

## Acknowledgments

This work was supported by the National Key R&D Program of China (2019YFA0906000, 2019YFA0906003), the National Natural Science Foundation of China (81772737, 31725002, 81972368). Guangdong Special Support Program (2021JC06Y578), Shenzhen Municipal Government of China (JCYJ20200109120016553, CJGJZD20200617102403009). Shenzhen High-Level Hospital Construction Fund and the Shenzhen Institute of Synthetic Biology Scientific Research Program (ZTXM20214005), The Sanming Project of Shenzhen Health and Family Planning Commission (SZSM201412018, SZSM201512037, SZSM202011017), Shenzhen Science and Technology Program (KQTD20180413181837372), Shenzhen Outstanding Talents Training Fund and Bureau of International Cooperation, Chinese Academy of Sciences (172644KYSB20180022). We thank the DNA assembly automation platform of China National GenBank for the support on synthetic chunk assembly. LAM and JSB were supported by NSF grants MCB-1026068, MCB-1443299, MCB-1616111 and MCB-1921641.

## Author contributions

Conceptualization, W.H., Y.S.; funding and resources, W.H., Y.S, Z.C.; microfluidic platform used for aging screening: Z.X.; data production, C.Z., Y.H., Y.A., D.Y, YL.W. data analyses, investigation, and visualization, Y.W., C.Z., X.F., J.Z.; writing – original draft, W.H., Y.S., C.Z, Y.W., and J.D.B.; writing – review & editing: all co-authors.

## Competing interests

Jef Boeke is a Founder and Director of CDI Labs, Inc., a Founder of and consultant to Neochromosome, Inc, a Founder, SAB member of and consultant to ReOpen Diagnostics, LLC and serves or served on the Scientific Advisory Board of the following: Sangamo, Inc., Logomix, Inc., Modern Meadow, Inc., Rome Therapeutics, Inc., Sample6, Inc., Tessera Therapeutics, Inc. and the Wyss Institute.

## STAR★METHODS

### RESOURCE AVAILABILITY

#### Lead contact

Further information and request for reagents and resources should be directed to and will be fulfilled by the lead contact.

#### Materials availability

All the requests for the generated plasmids and strains should be directed to the lead contact

#### Data and code availability

The data that support the findings of this study have been deposited into CNGB Sequence Archive of CNGBdb with accession number CNP0003780.

### EXPERIMENTAL MODEL AND SUBJECT DETAILS

#### Strains and growth media

The yeast strains used in this paper were derivatives of BY4741 or BY4742 ^29^. Synthesis of the minichunks was outsourced to BGI TECH SOLUTIONS (BEIJING LIUHE) CO., LIMITED. The strains generated in this study are listed in **Table S6**. Yeast culture and transformation were applied by using standard methods. The medium used in this study including two types: YPD and SC (synthetic complete medium) or synthetic medium without histidine (SC-His), synthetic medium without uracil (SC-Ura), synthetic medium without leucine (SC-Leu). Specifically, for replicative lifespan measurements, the cells were cultured on SC medium.

### METHOD DETAILS

#### *SynXIII* design and construction

Methods of synthetic chromosome design, synthesis, and construction described previously were used in this study^14,72^. The sequence of chromosome XIII was designed in silico and BioStudio following Sc2.0 design principles. The final version of designer chromosome XIII sequence (*synXIII*) was defined as yeast_chr13_3_40, with a total of 883,749 bp length and ~9% sequence modification, including removal of 21 tRNAs, insertion of 333 loxPsym sites, swap of TAG to TAA, and introduction of 529 pair synthetic PCRTags. More detailed information of *synXIII* can be accessed in Table S1).

The sequence of *synXIII* was hierarchically segmented to 15 megachunks (~70 kb, named A-O), then to 84 chunks (~ 10 kb), and final 421 minichunks (~3 kb) for commercial synthesis. For assembly verification, single colonies were selected for overnight culture at 37 °C. The restriction enzyme digestion was performed to verify the assembly result. An 800-1000 bp of the homologous region between each chunk was designed for homologous recombination and 40-bp overlap between each minichunk designed for Gibson assembly. In each megachunk integration, 5-6 chunks (equivalent to 1 megachunk) were directly transformed with the same moles of each chunk as a pool into yeast to replace the corresponding wild-type sequence. For synthetic chunks transformation, we adapted lithium acetate transformation strategy and each chunk was needed 300~500 ng which was excised from plasmids and purified by using AxyPrep (DNA Gel Extraction Kit) to screen the candidate colonies on the selective auxotroph medium plates. We integrated synthetic megachunks from both ends of chromosome XIII in parallel in strains BY4741 and BY4742. Nine successive rounds of left synthetic semi-arm integration and six rounds of right in the synthetic right-semi arm were used to produce the semi-synthetic synXIII strains (synXIII-I and synXIII-J, strain ID: Ycz011, Ycz017) which fused into full-length *synXIII* by I-*Sce*I mediated directed homologous recombination^22^. All strains generated in this study are listed in (**Table S6**).

#### Target colony selection

The colonies cultured on the selective plates were replicated onto the synthetic complete medium lacking uracil (SC–Ura) or leucine (SC–Leu). After overnight incubation at 30 °C, the clones can grow on one type of medium but not the other were identified and subjected to PCRtag analysis to verify the incorporation of the entire synthetic megachunk. To confirm the strains, two rounds of PCR were performed. At first, five or six pairs of PCRTags, distributed in each synthetic chunk, were chosen to screen the phenotypically desired clones. rTaq DNA polymerase (TaKaRa, R001W) was used together with 300 ng of genomic DNA in a 10 mL reaction containing 1 mM of primer each. The PCR program was as following:1 cycle of 94 °C for 5 min,35 cycles of 94 °C / 30 s-55 °C / 30 s-72 °C / 30 s, 1 cycle of 72 °C for 5 min and 12 °C keep. The clones got through the first round of PCR test were subjected to next round of PCRTag analysis using all primers within the megachunk to identify the ones containing the entire synthetic DNA.

#### Scanning electron microscope

The samples were grown in OD=0.1 for 4 hours to an A600 of 0.6, and then fixed in 2.5% glutaraldehyde (Sigma, catalog number: 340855, pH 7.4) for 2h. After washing three times with 0.1 M phosphate buffer (pH 7.2) and fixation in 1% osmic acid (Sigma, catalog number: 104239) at 4 °C for 2h, they were dehydrated through an ascending series of ethanol by ending in 100%, and then dried using Critical Point Dryer (Quorum, K850). Samples were coated with a very thin film of gold for 30 s using sputter coater (Cresstington, 108Auto). Samples were observed using an ultra-high resolution scanning electron microscope (Quasi-S, HITACHI Regulus 8100).

#### Immunoblot analysis

A single colony was picked and placed into 5 ml YPD liquid medium for overnight and next diluted into a total 5 ml fresh culture to A_600_ = 0.1. Until culturing at 30 °C for another 5 h, the cells were collected. Using 1 mL sterile water to resuspend the cells and the suspension was transferred to a new 1.5 mL EP tube, 10,000 g centrifugation for 1 min to collect the cells. 50 μL sterile water and 50 μL 0.2 M NaOH were added and maintained at room temperature for 5 min, followed by centrifugation for collected cells. Thereafter, 50 μL SDS sample buffer with 1% Triton X-20 was added, followed by heating to 95 °C for 5 min. The mixture was then centrifuged at 13,000 g for 10 min and the supernatant was aliquoted as the total protein extraction. 15 μL per lane for 10-lanes gel was used to detect Flag-*RRN9* and 2 μL per lane for 10-lanes gel to detect H3.

#### ChIP-qPCR

The strains were cultured in YPD medium at 30°C overnight. The culture as diluted with fresh YPD medium to A_600_ = 0.1 and further grown for 6 h with shaking until the A_600_= 0.8–1.0. Then the Formaldehyde (1% final) was used to crosslink at 25 °C for 25 min. Next, Glycine (0.15 M final) was added to abort the crosslink. ~50 A_600_ of cells were collected. After breaking the cells by glass beads and the chromatin sonicated by 0.5 μL micrococcal nuclease (MNase), 1/10 of the sonicated chromatin was took as the input and 9/10 were incubated with 2 μL anti-flag antibody for above 8 h. The antibody-protein-DNA complex was put down by Protein A/G Magnetic beads. To reverse the protein-DNA crosslinks 2.5 μL 10 mg/mL Rnase A and 5 μL 20 mg/mL Proteinase K was added and incubated at 65°C for another 8 h^73^. The DNA was purified by the kit DNA Clean & Concentrator-5 (ZYMO, catalog number: D4004). RT-PCR was performed to calculate the protein occupancy. All the occupancy data were shown as the percentage of enrichment at target loci normalized by *RPO21*^34^. For RT-PCR analysis, 50 μL of input samples were 10-diluted with TE as qPCR input samples (input versus chip dilution ratio was 100:1). RT-qPCR analyses were done with TB Green qPCR Kits (TAKARA, catalog number: AK8901). Each reaction with input or chip sample was performed under the following cycling condition: 95 ℃ 30 s and 39 cycles of 95 ℃ 5 s, 55 ℃ 10 s and 72 ℃ 30 s. Relative enrichments of ChIP versus input were quantified as input (N) = 2ΔCt (target gene)/2ΔCt (control gene). ChIP enrichment of each biological replicate (N = 3) was calculated from mean value of triplicated qPCR reactions. Primer sequences: *RRN9* forward primer 5’-gcaataccttccggtatat-3’, reverse primer 5’-caaccgctgataaagagac; *ACT1* forward primer 5’-atggattctgaggttgctgct-3’, reverse primer 5’-tggtgtcttggtctaccgac.

#### Mating type Switch of Yeast and Sporulation

PJD147 (pGal-HO) was transferred into the synXIII strain using the LiOAc transformation method and colonies were selected on SC–Leu plates. The colony was picked into SC–Leu (2% glucose), medium cultured overnight at 30 °C, then diluted reached A_600_= 0.1 in SC–Leu (2% galactose) fresh medium, followed by culturing for 4 hours, and then plating on YPD. The mating type was then confirmed by crossing *MAT***a** and *MAT*α tester strains (Ycz050 and Ycz051) and examined by microscopy. A strain with *MAT***a** and another strain with *MAT*α were recovered on YPD plates. The colonies from two strains were picked into YPD medium, cultured for 24 h and patched on YPD BYNplates. Diploids verified by testing with the two mating type testers were cultured and cells were collected and washed with sterile water for four times, following sporulated in SPOR medium containing 1% potassium acetate (Sigma, catalog number: 60035) and 0.125% yeast extract (Oxoid, catalog number: LP0021B) for three days.

#### Replicative lifespan measurement using microfluidic assay

A microfluidic device based on published reports was used to monitor the number of mother cell divisions.^39,74^ SC (synthetic complete) medium was used to grow all tested yeast strains. Cells incubated in 5 mL SC medium cultured overnight were diluted to an A_600_ of 0.1 in fresh medium, and incubated at 30 °C for 4~6 h until they reached an A_600_ of 0.6. The microfluidic device contains 8 micro-posts (side length in the range of 40-100 μm, depth of the chamber ranging between 3.8-4.2 μm) that clamp mother cells in place. In contrast, newly formed daughter cells are washed away by hydrodynamically controlled flow of the surrounding liquid medium. Each micro-post tested one strain combined with time-lapse microscopy at the high temporal solution. Mother cells were continuously monitored for 60 h repeated microscopic imaging to measure produced daughter cells (**Figure S5A**).^39^ For each strain, 40 mother cells were randomly selected, the number of daughter cells produced was detected, and the survival curve was drawn using Matlab.

#### SCRaMbLEant library preparation

An enhanced green fluorescent gene (*eGFP*) was inserted before the stop codon of *HSP104* fused with *LEU2* marker for selection. pSCW11-*Cre/EBD-HIS3* was transferred into the strain and simultaneously a *URA3* marker was integrated into the middle of two loxPsym sites between *YMR011W* and *YMR012W* on *synXIII* at position (270,117 ~ 270,525 bp). The colony was cultured in the SC–His medium overnight and then diluted to an A_600_ of 0.1 in the 5 mL fresh SC–His medium containing the inducer 1 µM estradiol at 200 rpm in a shaking flask, with induction for 8 h at 30 °C.

Finally, 200 μL solution was collected and the cells were washed by sterile water, culturing on a SC+5-FOA plates for 3 days. For each plate, 96 FOA-resistant single colonies were isolated from the plate for culturing with SC liquid medium in 96 deep-well plate. Then overnight culture for all plates were prepared for GFP fluorescence measurement. In this way, a pool of about 6000 SCRaMbLEant colonies was produced.

#### FACS testing

Each SCRaMbLEant colony was grown in 1 mL SC medium of a 2.2 mL deep well plate for 24 h. The following day, cells were diluted to an A_600_ of 0.001 in 1 mL fresh medium and grown up to an A_600_ of 0.8, and each strain was tested by flow cytometry (BD, FACSAria Cell Sorter; Beckman, CytoFLEX). Analysis and cell sorting with flow cytometry were conducted at a rate of 10,000 events per second. The 488 nm laser was used to excite the GFP protein and 528/29 filter was used to detect the fluorescence signal. An initial scatter-gating step was operated based on cell’s forward-scatter properties to collect data from single cells. Flow cytometry analysis was performed by analyzing 100,000 cells of each sample.

#### Omics analyses

Whole-genome sequencing was performed for the synXIII (yeast_chr13_9_2, strain ID: Yzc020) on the NextSeq 500 platform. For transcriptome and proteome, yeast cells Yzc030 and BY4741, with three biological replicates of each were cultured and collected until an A600 of ~ 0.8 was reached. The RNA sequencing libraries were constructed using NEB Next Ultra TMRNA Library Prep Kit (catalog number: E7770S, NEB). Then sequencing was performed on Illumina HiSeq PE150 Sequencing Systems by v2 kit (catalog number: 256782). For proteome, a free-labeled peptide method described previously was used in this study. The peptides were loaded on an EASY-nLCTM 1200 UHPLC by the autosampler onto a 2 cm C18 trap column and an analytical C18 column packed in-house (0.075 × 150 mm column, 3 μm). The peptides were subjected to nano-electrospray ionization (nano ESI) followed by tandem mass spectrometry (MS/MS) in a benchtop Orbitrap Q Exactive mass spectrometer (Thermo Fisher Scientific, San Jose, CA) coupled online tSYNSYYo the HPLC. The method of omics data processing described previously was used in this study^22^. For nucleotide sequence analysis, the sequencing data quality control was performed using SOAPnuke; the clean data were aligned to reference sequence of synXIII yeast genome using Bowtie2 (version 2.2.5).^75^ And the variations were identified with both GATK (version 2.7)^76^ and SAMtools (version 0.1.19).^77^ For transcriptomics, the reads were mapped to genomes by hisat2 (version 2.1.0)^78^, and the quantification and differential expression genes were analyzed by HTSeq featureCounts (version 2.0.1)^79^ and DEseq2(version v1.30.1).^80^ For proteomics, Mascot (version: 2.8.0) and IQuant (version: 2.0.1) were used for protein identification and quantification.^81^ False positive differentially expressed genes were inferred by the following rules and removed: dubious genes, transposable genes, 2-micron genes or genes with low coverage (<60%) for native and synthetic strains. Gene enrichment of KEGG pathways and Gene Ontology annotations were performed using the hyper-geometric and Chi-squared tests.

#### WGCNA analysis and hub gene identification

The weighted co-expression networks of SCRaMbLEd yeast strains were constructed using the WGCNA R package (version 1.69).^82,83^ The normalized TPM (Transcripts Per Kilobase Million) of each gene was considered as input data. The soft-thresholding power of nine was used to construct the scale-free network. The module eigengene (MEs, co-expression gene cluster) were selected using a dynamic tree cut method with a threshold of 0.35 and the minimum number of genes in each module at 30. The correlations between modules and phenotype traits of strains were investigated using the Pearson correlation. The co-expression network was displayed by Cytoscape software (version 3.8.0).^84^ The core objects with high connectivity in scale-free networks were identified by Maximal Clique Centrality (MCC) method using cytoHubba (version 1.0.1).^85^ These core objects with a correlation coefficient of gene significance (GS) and module membership (MM) >= 0.8 were identified as final core genes, highly related to the phenotypes

#### Protein-Protein Interaction (PPI) network and target gene identification

The PPI networks were constructed using Cytoscape software (version 3.8.0).^84^ The target genes were analyzed by inputting them into the Search Tool for the Retrieval of Interacting Genes (STRING version 11.0). A confidence score ≥ 0.3 was used to construct a protein-protein interaction (PPI) network. The network layout was calculated by Edge-weighted Spring Embedded Layout Algorithm in Cytoscape. The biological function of sub network was identified by STRING enrichment.

### Quantification and statistical analysis

The data was analyzed using GraphPad Prism (version 9.4.0). One-tailed t-tests were used to compare different groups in this paper. Error bars represent SD. Differences were considered as statistically significant at p-valueL<L0.05. * represents PL<L0.05, ** represents PL<L0.01, *** represents PL<L0.001, **** represents PL<L0.0001

---

**Figure S1.**
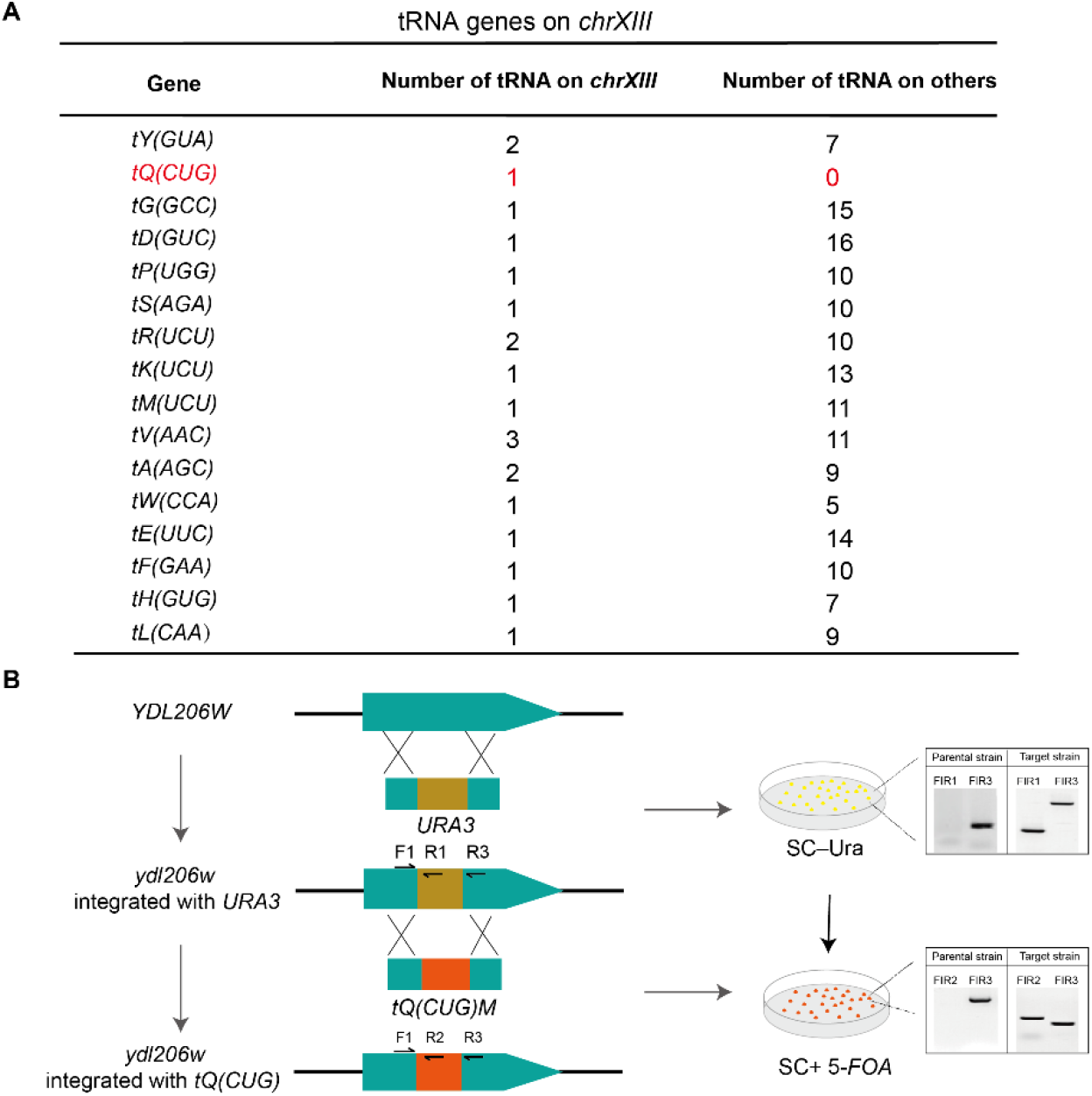
Summary of tRNAs in *chrXIII*. **(A)** Summary of tRNA in *chrXIII* and compared with other 15 chromosomes. (**B**) The design of *tQ(CUG)* relocation into *YDL206W*.

**Figure S2.**
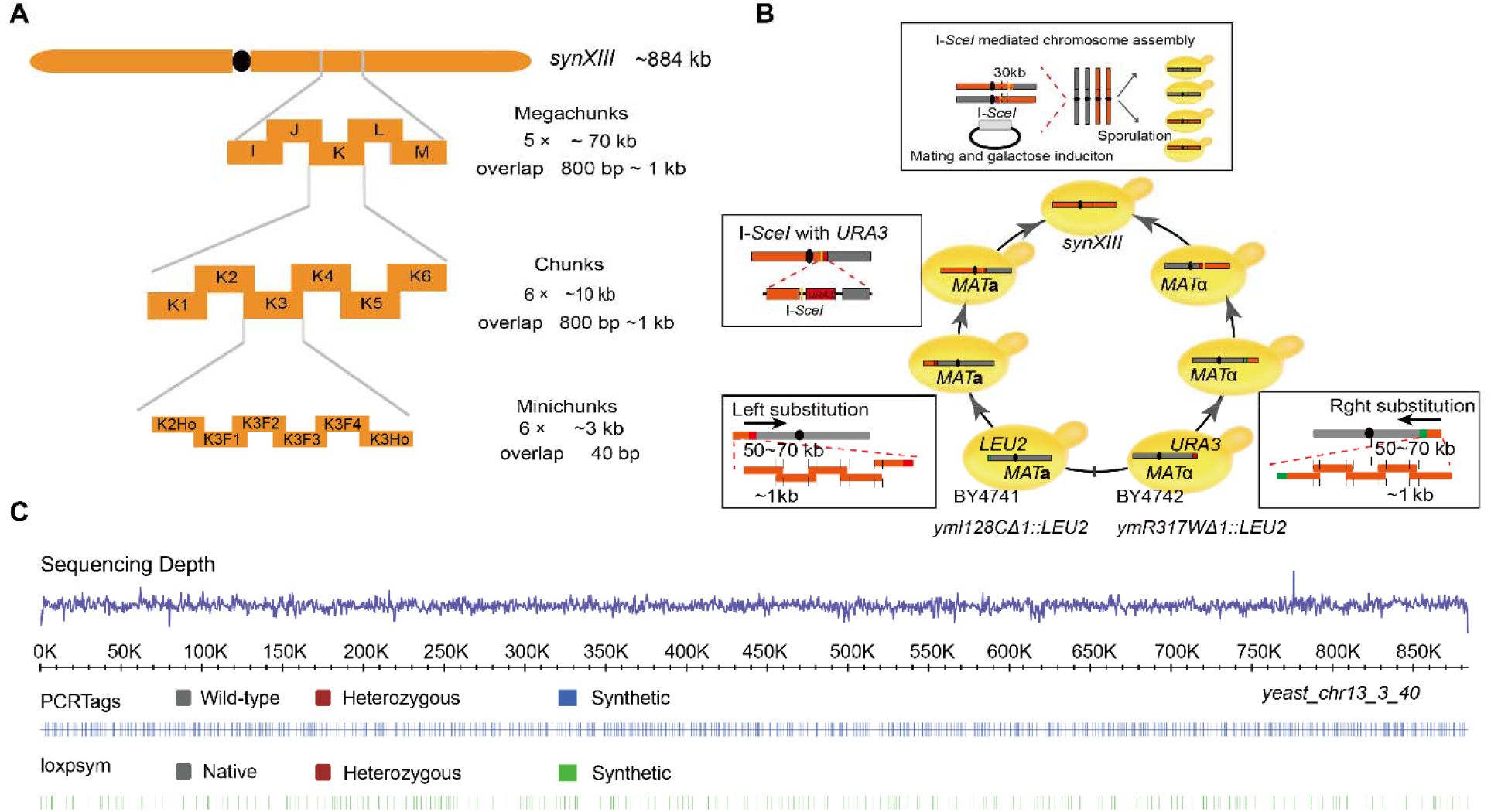
Synthetic chromosome XIII construction. **(A)** The intact *synXIII* chromosome are segmented stepwise into ~50 kb megachunks (as K, L, M), further divided into ~10kb chunks (such as K1, K2, K3) and further divided into ~3 kb minichunks (such as K1FI, K1F2) according to the experimental design. More details are provided in Supplementary text. (**B**) Native chromosome XIII replacement with synthetic chunks. On average 5 chunks were used to replace the native segments of chromosome XIII (dark grey). 9 and 6 step-by-step replacements were carried out in parallel from the left arm and right arm respectively with iterative selectable markers (LEU2 or URA3) through homologous recombination in yeast with homologous regions around 500 bp. ~1 kb junctions (purple) produced by PCR using ligated chunks as template was designed to have 500 bp overlapping regions with two adjacent chunks to improve the replacement efficiency. I-*Sce*I mediated synXIIIA-I and synXIIIJ-O integration. An I-*Sce*I site (yellow) was introduced on both semi-synthetic chromosomes. After mating, I-*Sce*I induction was performed to generate double strain breaks on both semi-synthetic chromosomes. 30 kb homologous region on both semi synthetic chromosomes enabled the integration of complete *synXIII* (blue). (**C**) Whole genome sequencing analysis of synthetic chromosome XIII. The sequencing depth indicates its average in 500-bp windows. The PCRtags and loxPsym site were identified by the sequencing reads containing its synthetic-specific sequence (supporting read ≥ 5).

**Figure S3.**
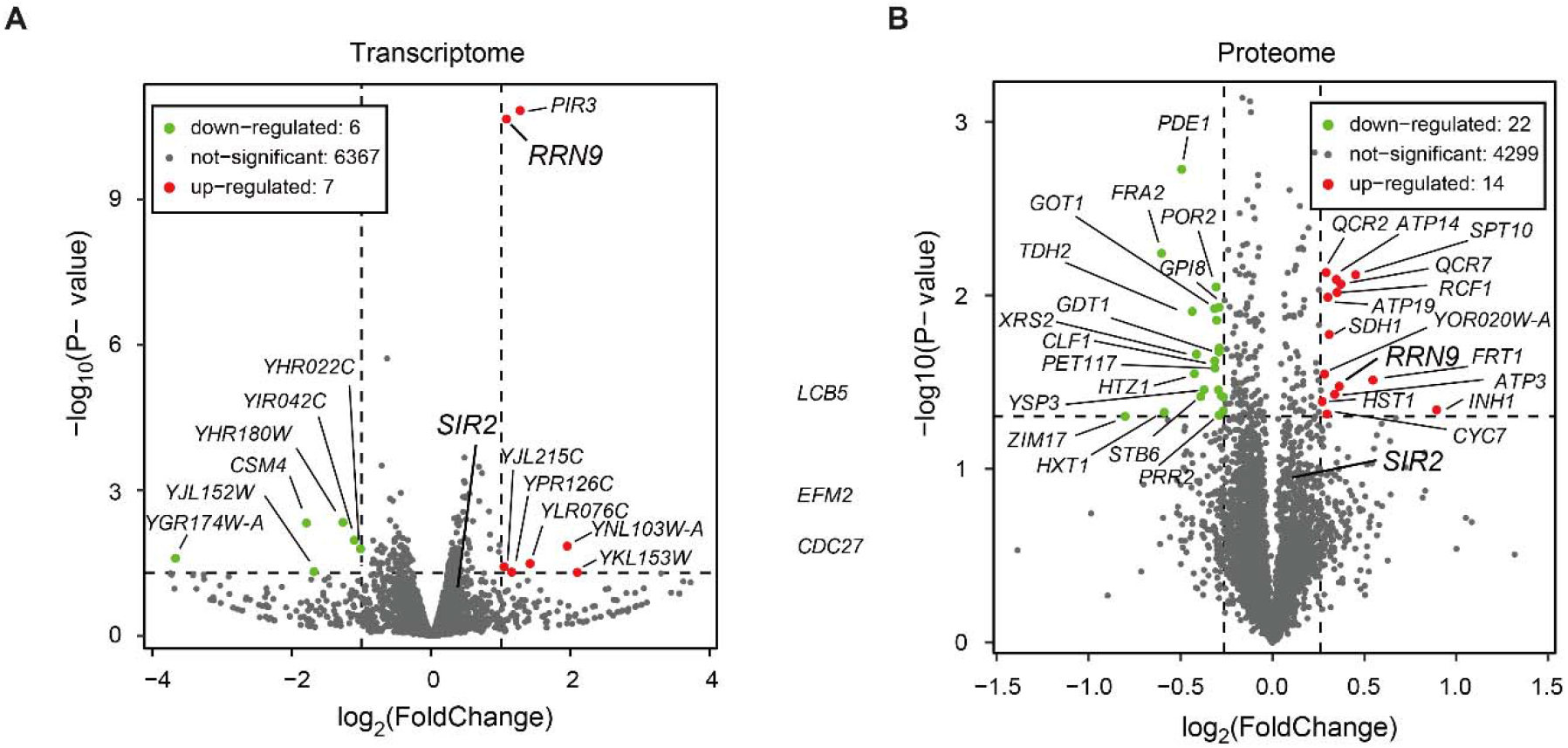
The transcriptome and proteome analysis of Ycz403 strain in comparison with Ycz402. **(A)** Transcriptomic and **(B)** proteomic level of gene differential expression in Ycz403 compared to Ycz402. The up and down-regulated genes are labeled in red and green respectively.

**Figure S4.**
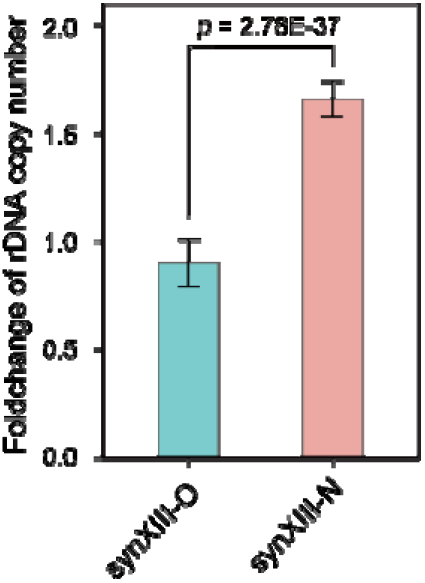
The foldchange of rDNA copy number of synXIII-N compared to synXIII-O. The rDNA copy number calculated by an average of sequencing depth in each 500-bp window normalized to the sequencing depth of whole genome. The statistical confidence is calculated by one-sided T-test, and error bars represent standard division.

**Figure S5.**
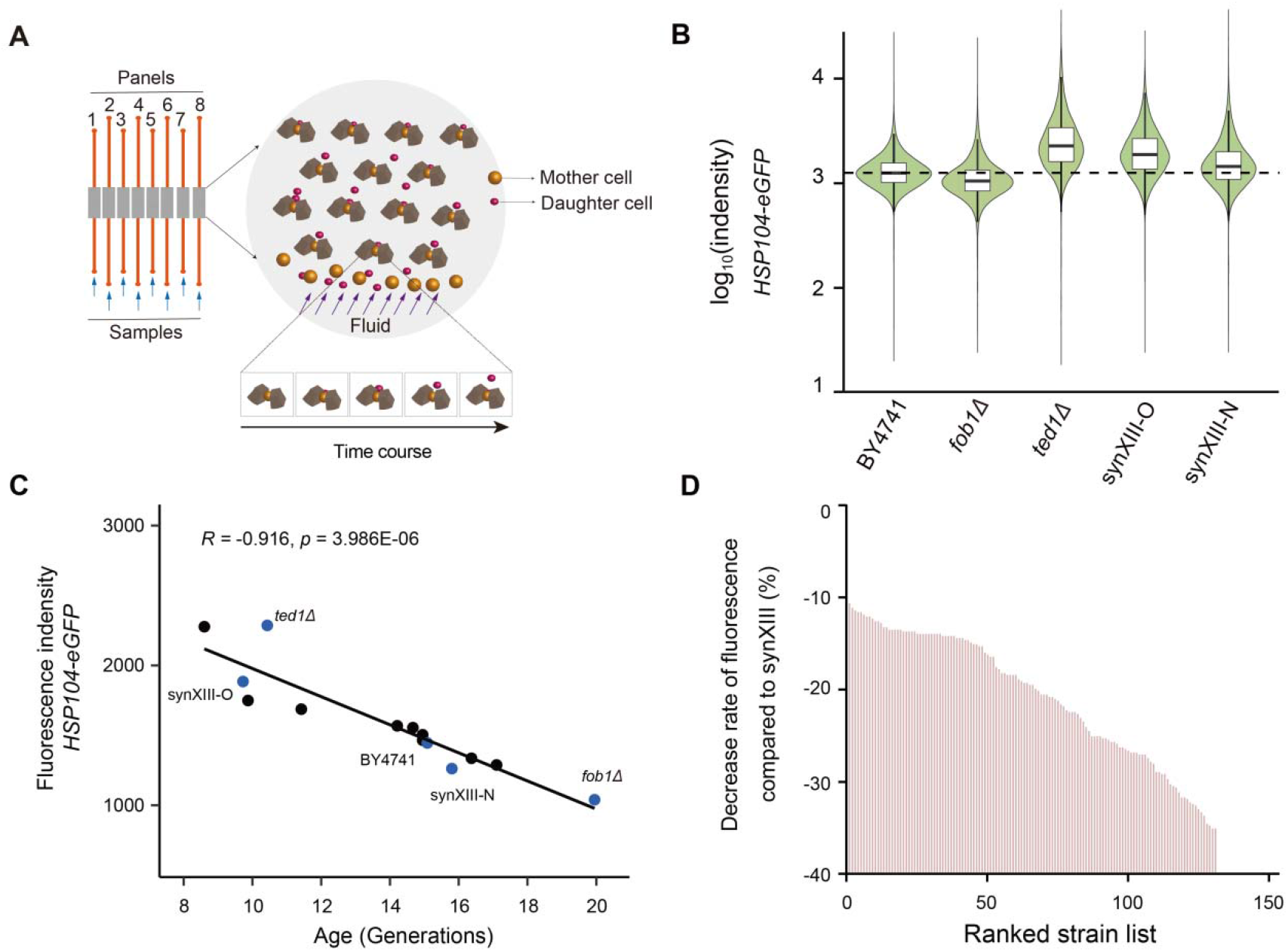
Lifespan measurement using *HSP104* as an aging biomarker. **(A)** Schematic for microfluidic system for automating lifespan measurement. Each chip with 8 micro-channels (left) is sealed on a thin glass slide (bottom). Geometries of the channels (red) and chambers (brown). Arrows show the directions of fluid flow, the mother cell with the bigger size stuck by chamber and the daughter cell washed away by fluid. The divisions of mother cell represent its replicative lifespan. (**B)** Comparison of *HSP104-eGFP* expression for BY4741, *ted1*Δ, *fob1*Δ, synXIII-O and synXIII-N. The *ted1*Δ and *fob1*Δ strains were constructed by using *URA3* to replace the ORF region of *TED1* and *FOB1*. The fluorescence intensity of *HSP104-eGFP* was measured by FACS (~100,000 cells for each sample). (**C)** The correlation between *HSP104-eGFP* and the replicative lifespan across genetic variant mutants. Each measurement with three replicates and 40 cells were monitored for each replicate of lifespan analysis. **(D)** The decreased rate of fluorescence intensity comparing to synXIII. The fluorescence intensity of *HSP104-eGFP* was measured by FACS (~100,000 cells for each sample).

**Figure S6.**
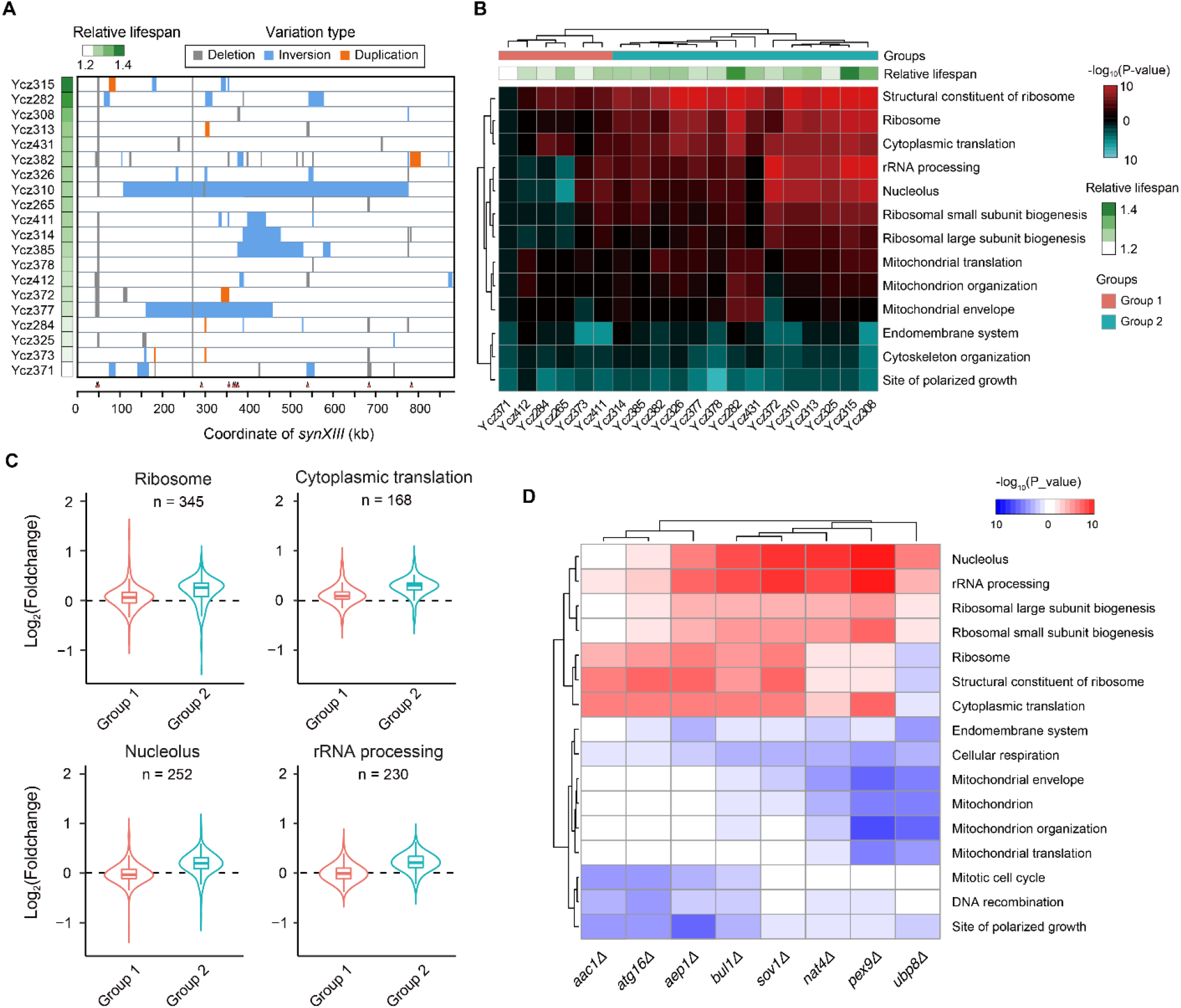
Function enrichment analysis of 20 lifespan-extended SCRaMbLEants. **(A)** Schematic map of structural variations in 20 long-lived SCRaMbLEants. Y-axis shows the relative aging extension of each SCRaMbLEant in comparison to parental synXIII strain that of represented by color scale. The red rectangles indicate the loci of aging regulators on chromosome *synXIII*. **(B)** Transcriptome profiling of lifespan extended strains compared to synXIII strain Ycz063. The heatmap represents the statistical significance of Gene Ontology (GO) enrichment of differentially expressed genes. red, up-regulated; green, down-regulated. Long-lived strains were clustered into three groups by hierarchical clustering, named “Group 1” and “Group 2”. **(C)** The differential expression analysis of genes involving in significantly altered GO categories. The expression level of genes represented by the median of strains in each group. The number indicated the scale of genes for each GO category. (D) Transcriptome profiling of null mutant strain compared to BY4741. The heatmap represents the statistical significance of Gene Ontology (GO) enrichment of differentially expressed genes; red, up-regulated; blue, down-regulated.

**Table S1.** Summary of *synXIII* design.

| Modification | Design feature | Number | Base alteration |
| --- | --- | --- | --- |
| Replacement | wild type telomere -> Universal<br>Telomere Cap<sup>a</sup> | 2 | 16880->1378 |
|  | 629 pairs of WT PCR tags and 629 pairs<br>of SYN PCRTags | 1258 | 15879 |
|  | stop codon TAG -> TAA | 100 | 100 |
| Deletion | transposable element region<sup>b</sup> | 36 | 28246 |
|  | gene | 21 | 30187 |
|  | tRNA | 21 | 1691 |
|  | snoRNA | 2 | 175 |
|  | intron<sup>c</sup> | 17 | 2707 |
| Insertion | LoxPsym site | 333 | 11322 |

**a.** Wild-type telomeres together with its adjacent subtelomeric regions are replaced with universal telomere caps. **b.** Transposable element region include these functional features of LTR retrotransposons, transposable element genes and long terminal repeats. **c.** Nine introns were retained as these introns reside in ribosomal subunit coding genes.

**Table S2.**
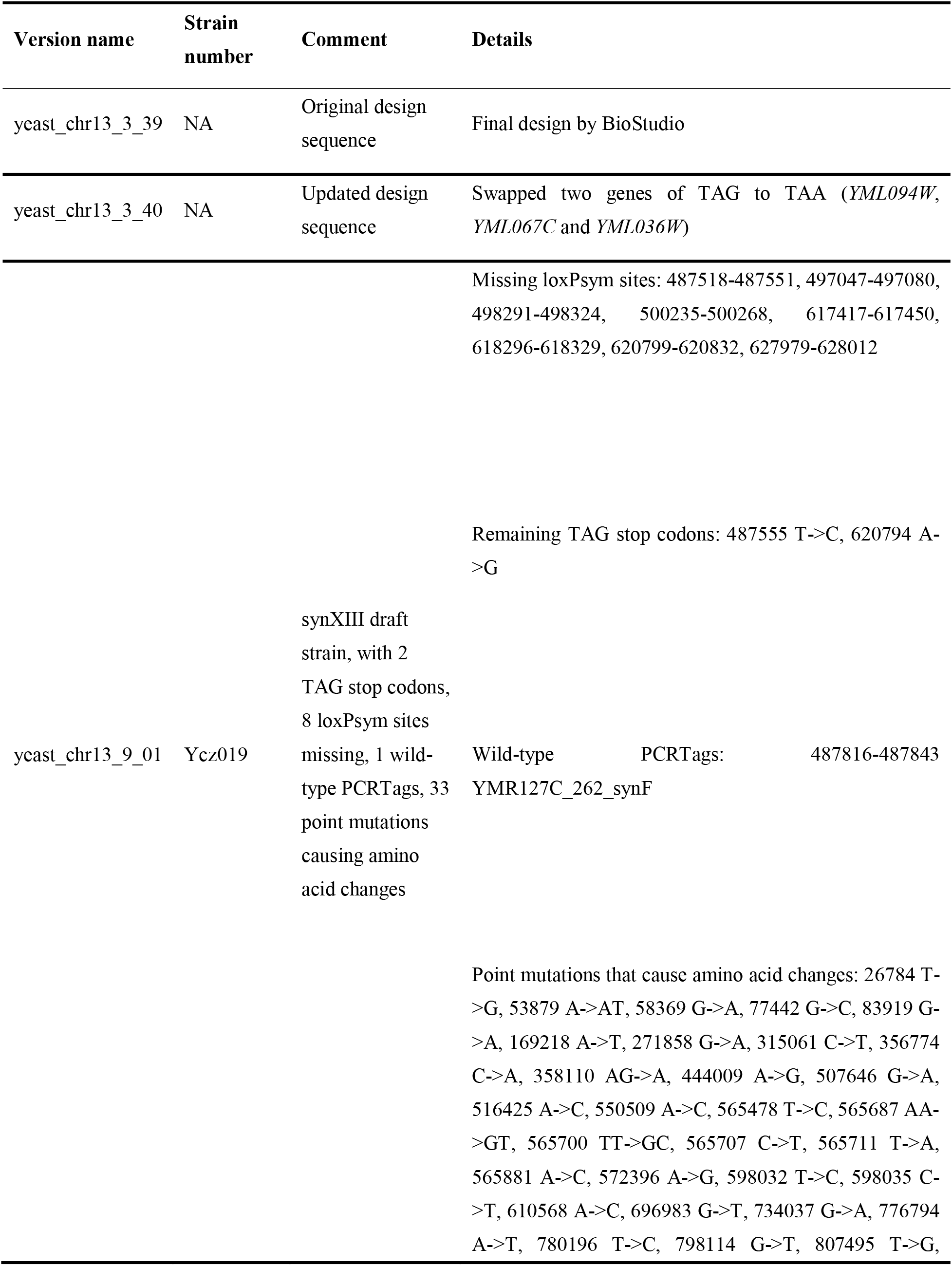
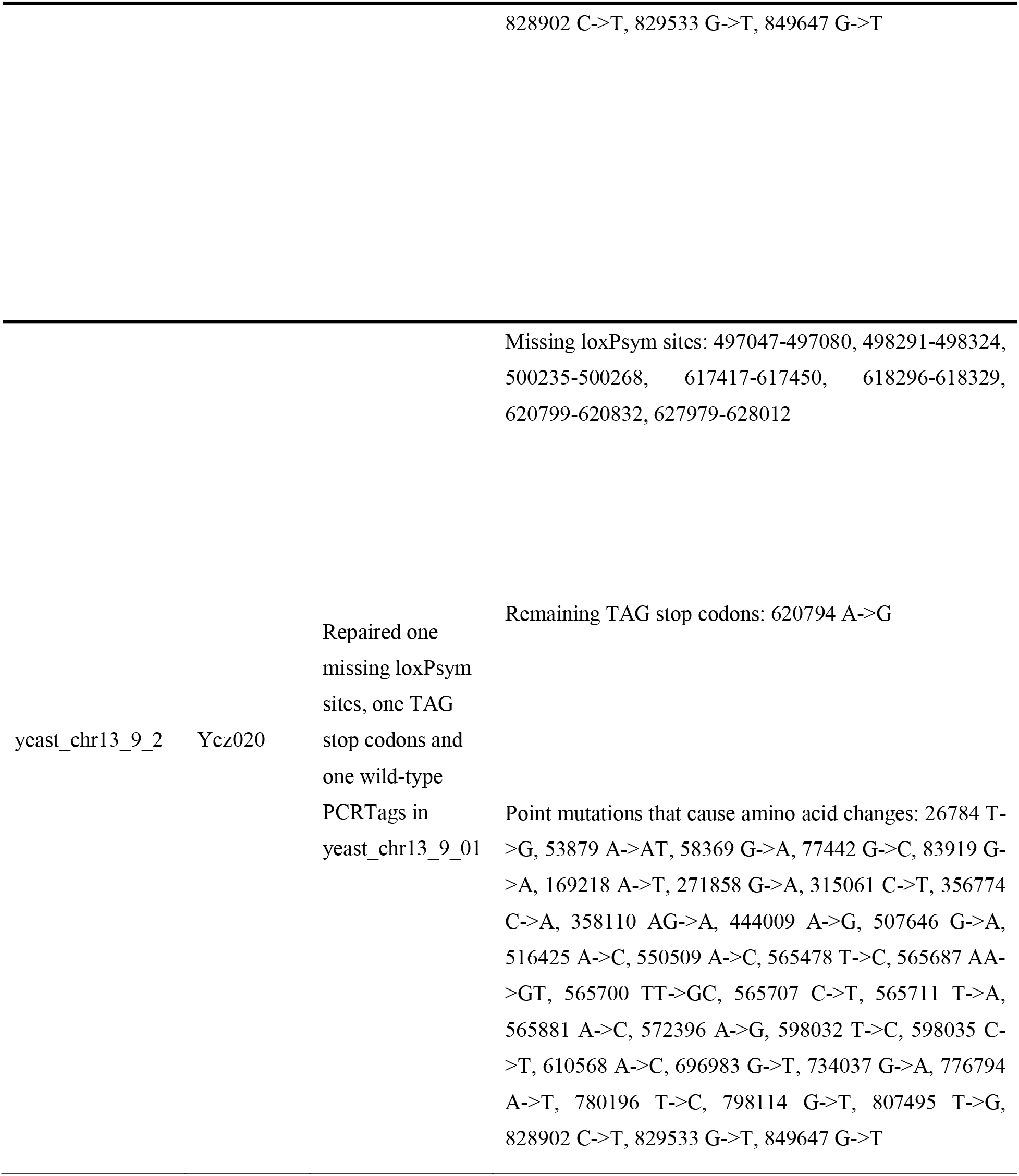
Chromosome version used in this paper.

**Table S3. Variant gene list from 135 SCRaMbLEd strains**

321 SCRaMbLEd genes with different type of changes are listed in the excel file of Table S3.

**Table S4. SCRaMbLE events of 20 long-lived strains**

SCRaMbLE events and its variant genes are listed in the excel file of Table S4.

**Table S5.** Aging regulators identified by WGCNA analysis.

| GeneID | ChrID | Gene<br>name | Essential<br>status | GS<sup>a</sup> | p-value of<br>GS | MM<sup>b</sup> | P-value of<br>MM | logMCC<sup>c</sup> |
| --- | --- | --- | --- | --- | --- | --- | --- | --- |
| YLR035C | chr12 | MLH2 | Nonessential | 0.801 | 2.22E-05 | 0.796 | 2.71E-05 | 13.96 |
| YDL220C | chr04 | CDC13* | Essential | 0.792 | 3.20E-05 | 0.857 | 1.38E-06 | 13.96 |
| YBR275C | chr02 | RIF1* | Nonessential | 0.765 | 8.50E-05 | 0.795 | 2.78E-05 | 10.14 |
| YJL194W | chr10 | DDC1* | Essential | 0.760 | 0.000101 | 0.737 | 0.0002073 | 13.96 |
| YNL126W | chr14 | SPC98 | Essential | 0.755 | 0.000120 | 0.720 | 0.000342 | 13.96 |
| YAR035W | chr01 | YAT1* | Nonessential | 0.747 | 0.000152 | 0.791 | 3.35E-05 | 8.72 |
| YDR027C | chr04 | VPS54* | fast_growth | 0.744 | 0.000167 | 0.863 | 9.89E-07 | 11.07 |
| YPL194W | chr16 | CDC6 | Nonessential | 0.740 | 0.000190 | 0.878 | 3.63E-07 | 13.96 |
| YAL002W | chr01 | VPS8* | Nonessential | 0.735 | 0.000221 | 0.794 | 2.95E-05 | 9.02 |
| YOR100C | chr15 | CRC1* | Nonessential | 0.717 | 0.000371 | 0.673 | 0.0011339 | 8.72 |

**a.** GS, gene significance indicates the correlation of its gene expression profile with replicative aging. **b.** MM, module membership presents the correlation of its gene expression profile with the module eigengene. **c.** MMC, The Maximal Clique Centrality of corresponding PPI network calculated by CytoHubba APP in Cytoscape. \* Indicated the genes are related to the aging in GenAge or SGD Database.

**Table S6. Yeast strains used in this paper**

All strains used in this paper listed in the excel file of Table S6.

## Notes

### Competing Interest Statement

The authors have declared no competing interest.

